# MYC-Driven Activation of USP39 Enhances SRSF1 Stability and Promotes PDAC Progression

**DOI:** 10.1101/2025.08.22.671882

**Authors:** Benteng Ma, Alexander J. Kral, Xin Zhang, Neelu Singh, Astrid Deschenes, Paolo Cifani, Youngkyu Park, David A. Tuveson, Adrian R. Krainer, Ledong Wan

## Abstract

Pancreatic ductal adenocarcinoma (PDAC) is a highly lethal malignancy primarily driven by oncogenic KRAS signaling. The splicing factor SRSF1 plays a key oncogenic role in PDAC, where its tightly regulated expression constrains KRAS-driven signaling under normal conditions, while its upregulation promotes tumorigenesis. SRSF1 expression is regulated in part by proteostasis. However, the precise mechanisms remain unclear. Here, we identify USP39 as a deubiquitinase that interacts with SRSF1 in an RNA-independent manner and stabilizes it by reducing ubiquitination. USP39 expression is elevated in PDAC tumors and precancerous lesions, and correlates with poor patient survival. *USP39* knockdown suppresses PDAC cell proliferation and migration, effects that are partially rescued by SRSF1 overexpression. Mechanistically, we show that MYC directly activates *USP39* transcription via E-box motifs within its exon 1b promoter, linking MYC-driven transcriptional regulation to post-translational stabilization of SRSF1. Together, these findings define a MYC-USP39-SRSF1 regulatory axis that integrates transcriptional and post-translational mechanisms in PDAC and highlight USP39 as a potential therapeutic target.

## INTRODUCTION

Pancreatic ductal adenocarcinoma (PDAC) is one of the deadliest malignancies, with a dismal 5-year survival rate of approximately 13%.^1^ Somatic activating mutations in KRAS, a small GTPase in the RAS superfamily, are present in the vast majority of PDAC cases. The most prevalent variant, KRAS^G12D^—a glycine-to-aspartate substitution at codon 12 within the GTP-binding domain—accounts for ∼40% of PDAC cases and plays a key role in initiating precancerous transformation.^2, 3^ Tumor progression is further driven by co-occurring inactivating mutations in tumor suppressor genes, such as *TP53*, *CDKN2A*, and *SMAD4*.^1, 4^

Post-translational regulation of protein stability by the ubiquitin–proteasome system (UPS) plays a critical role in maintaining cellular homeostasis.^5^ It is responsible for 80–90% of cellular proteolysis and 10–20% of autophagy.^6^ Ubiquitination involves the covalent attachment of ubiquitin to substrate proteins by ubiquitinases, while deubiquitinases (DUBs) counteract this process by removing ubiquitin chains.^6, 7^ Ubiquitylation and deubiquitylation play essential roles in regulating protein stability, localization, and metabolism, as well as in controlling cellular physiological and pathological processes.^6–10^ Recent studies have implicated dysregulated ubiquitination and deubiquitination in multiple cancer-related pathways, including cell-cycle progression, metabolic rewiring, and immune evasion.^11, 12^

Dysregulated expression or mutations of oncogenic splicing factors can contribute to tumor development by altering pre-mRNA alternative splicing events that impinge on multiple signaling pathways.^13–16^ Serine/arginine-rich splicing factor 1 (SRSF1) is a highly conserved splicing factor that recognizes specific RNA sequences and regulates alternative splicing.^17–19^ We previously reported that SRSF1 is subject to ubiquitin-mediated destabilization in phenotypically normal pancreatic epithelial cells expressing KRAS^G12D^, forming a negative feedback loop that restrains MAPK signaling and maintains epithelial homeostasis.^19^ This regulation is overridden by MYC overexpression, which enhances SRSF1 transcription.^19^ SRSF1 expression is regulated through multiple mechanisms, including transcriptional control, alternative splicing, translational regulation, and protein stability.^20–25^ We also reported that MYC activation attenuates the ubiquitination-mediated regulation in mouse pancreas.^19^ However, the functional significance of SRSF1 ubiquitination in promoting its oncogenic activity remains to be elucidated.

Here we report that SRSF1 interacts with the DUB USP39 in an RNA-independent manner. USP39 stabilizes SRSF1 protein by decreasing its ubiquitination. Using multiple tumor databases and clinical PDAC samples, we demonstrate that USP39 functions as an oncogene and is associated with poor patient survival. Moreover, we show that MYC binds to the *USP39* promoter and enhances its transcription, thereby promoting SRSF1 expression through post-translational mechanisms.

## RESULTS

### SRSF1 interacts specifically with USP39

To identify SRSF1 interactors associated with the ubiquitin–proteasome pathway, we performed immunoprecipitation (IP) followed by mass spectrometry (MS) in SUIT-2 PDAC cells expressing T7-tagged SRSF1. This analysis identified eight proteins associated with the ubiquitin–proteasome pathway that were enriched in the SRSF1 pulldown (Fig. 1A). In parallel, we reanalyzed our previously reported SRSF1 interactome generated by proximity labeling in HeLa cells,^26^ which revealed seventeen proteins involved in the ubiquitin–proteasome pathway in close proximity to SRSF1 (Fig. 1A, Table 1). Four proteins—USP39, PSMD7, PSMD1, and CUL4B—were consistently detected in both experiments. These proteins were robustly enriched in proximity labeling using BioID2 fused to either the N- or C-terminus of SRSF1, at levels comparable to SRPK1 (Fig. 1B)—a known SRSF1-binding kinase that phosphorylates SRSF1^27,28^.

**Figure 1.**
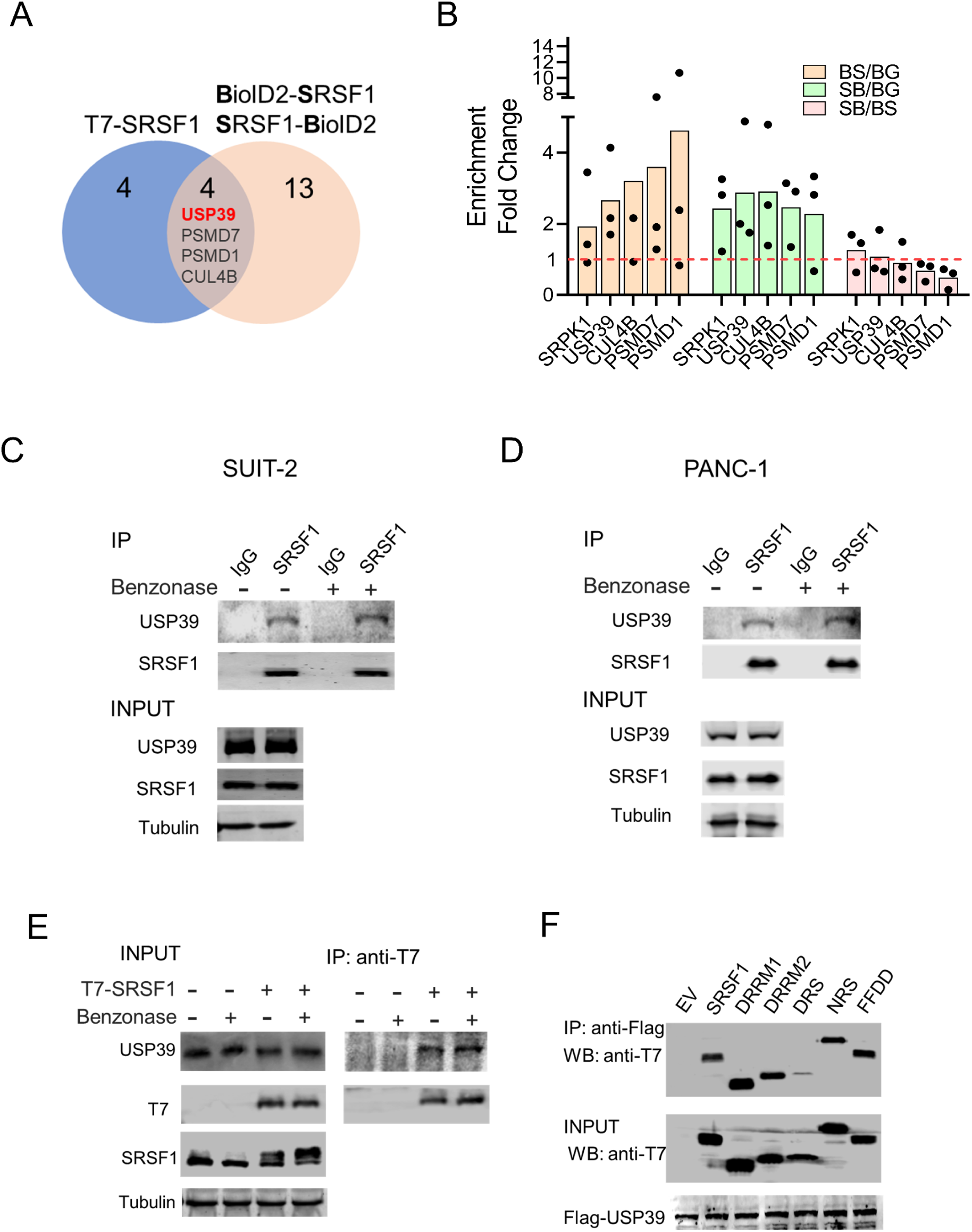
SRSF1 interacts specifically with USP39. (A) Venn diagram showing the overlap of ubiquitin–proteasome pathway-associated proteins identified by SRSF1 immunoprecipitation in SUIT-2 T7-SRSF1 cells and by proximity labeling in HeLa cells using BioID2 fused to either the N- or C-terminus of SRSF1^26^. (B) Relative protein enrichment in the proximity-labeling datasets compared to the BioID2-GFP control (n=3). BS: BioID2-SRSF1; SB: SRSF1-BioID2; BG: BioID2- EGFP. The known interactor, SRPK1, is included as a control. (C-D) SRSF1 immunoprecipitation in SUIT-2 (C) and PANC-1 (D) cells, with or without the addition of Benzonase to the cell lysate. Co-immunoprecipitated proteins were detected by immunoblotting with the indicated antibodies. (E) Co-immunoprecipitation assays were performed in SUIT-2 cells expressing T7-SRSF1, with or without the addition of Benzonase to the cell lysate. (F) Co-immunoprecipitation assays were performed in 293T cells expressing Flag-USP39 along with domain-deleted or mutated T7- SRSF1. SRSF1 deletion mutants lack either RRM1 (DRRM1), RRM2 (DRRM2), or the RS domain (DRS); the inactivated RRM1 mutant of SRSF1 (FFDD) has Phe56 and Phe58 substituted with Asp residues.^40^ NRS consists of a C-terminal fusion of SRSF1 with the nuclear retention signal from SRSF2.^41^

**Table 1.**
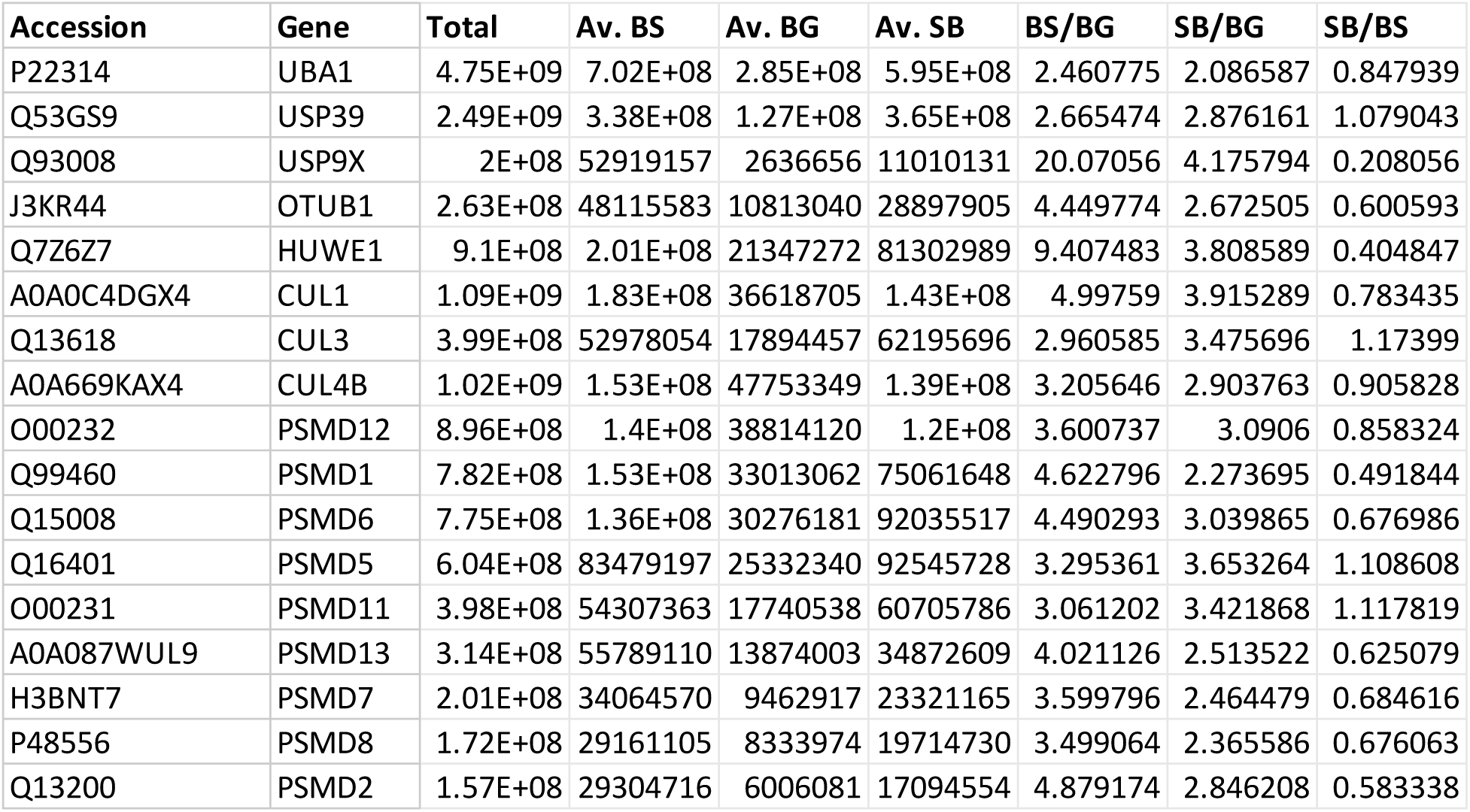
Proteins associated with the ubiquitin–proteasome pathway identified by proximity labeling of SRSF1.

| Accession | Gene | Total | Av. BS | Av. BG | Av. SB | BS/BG | SB/BG | SB/BS |
| --- | --- | --- | --- | --- | --- | --- | --- | --- |
| P22314 | UBA1 | 4.75E+09 | 7.02E+08 | 2.85E+08 | 5.95E+08 | 2.460775 | 2.086587 | 0.847939 |
| Q53GS9 | USP39 | 2.49E+09 | 3.38E+08 | 1.27E+08 | 3.65E+08 | 2.665474 | 2.876161 | 1.079043 |
| Q93008 | USP9X | 2E+08 | 52919157 | 2636656 | 11010131 | 20.07056 | 4.175794 | 0.208056 |
| J3KR44 | OTUB1 | 2.63E+08 | 48115583 | 10813040 | 28897905 | 4.449774 | 2.672505 | 0.600593 |
| Q7Z6Z7 | HUWE1 | 9.1E+08 | 2.01E+08 | 21347272 | 81302989 | 9.407483 | 3.808589 | 0.404847 |
| A0A0C4DGX4 | CUL1 | 1.09E+09 | 1.83E+08 | 36618705 | 1.43E+08 | 4.99759 | 3.915289 | 0.783435 |
| Q13618 | CUL3 | 3.99E+08 | 52978054 | 17894457 | 62195696 | 2.960585 | 3.475696 | 1.17399 |
| A0A669KAX4 | CUL4B | 1.02E+09 | 1.53E+08 | 47753349 | 1.39E+08 | 3.205646 | 2.903763 | 0.905828 |
| O00232 | PSMD12 | 8.96E+08 | 1.4E+08 | 38814120 | 1.2E+08 | 3.600737 | 3.0906 | 0.858324 |
| Q99460 | PSMD1 | 7.82E+08 | 1.53E+08 | 33013062 | 75061648 | 4.622796 | 2.273695 | 0.491844 |
| Q15008 | PSMD6 | 7.75E+08 | 1.36E+08 | 30276181 | 92035517 | 4.490293 | 3.039865 | 0.676986 |
| Q16401 | PSMD5 | 6.04E+08 | 83479197 | 25332340 | 92545728 | 3.295361 | 3.653264 | 1.108608 |
| O00231 | PSMD11 | 3.98E+08 | 54307363 | 17740538 | 60705786 | 3.061202 | 3.421868 | 1.117819 |
| A0A087WUL9 | PSMD13 | 3.14E+08 | 55789110 | 13874003 | 34872609 | 4.021126 | 2.513522 | 0.625079 |
| H3BNT7 | PSMD7 | 2.01E+08 | 34064570 | 9462917 | 23321165 | 3.599796 | 2.464479 | 0.684616 |
| P48556 | PSMD8 | 1.72E+08 | 29161105 | 8333974 | 19714730 | 3.499064 | 2.365586 | 0.676063 |
| Q13200 | PSMD2 | 1.57E+08 | 29304716 | 6006081 | 17094554 | 4.879174 | 2.846208 | 0.583338 |

**Table 2.**
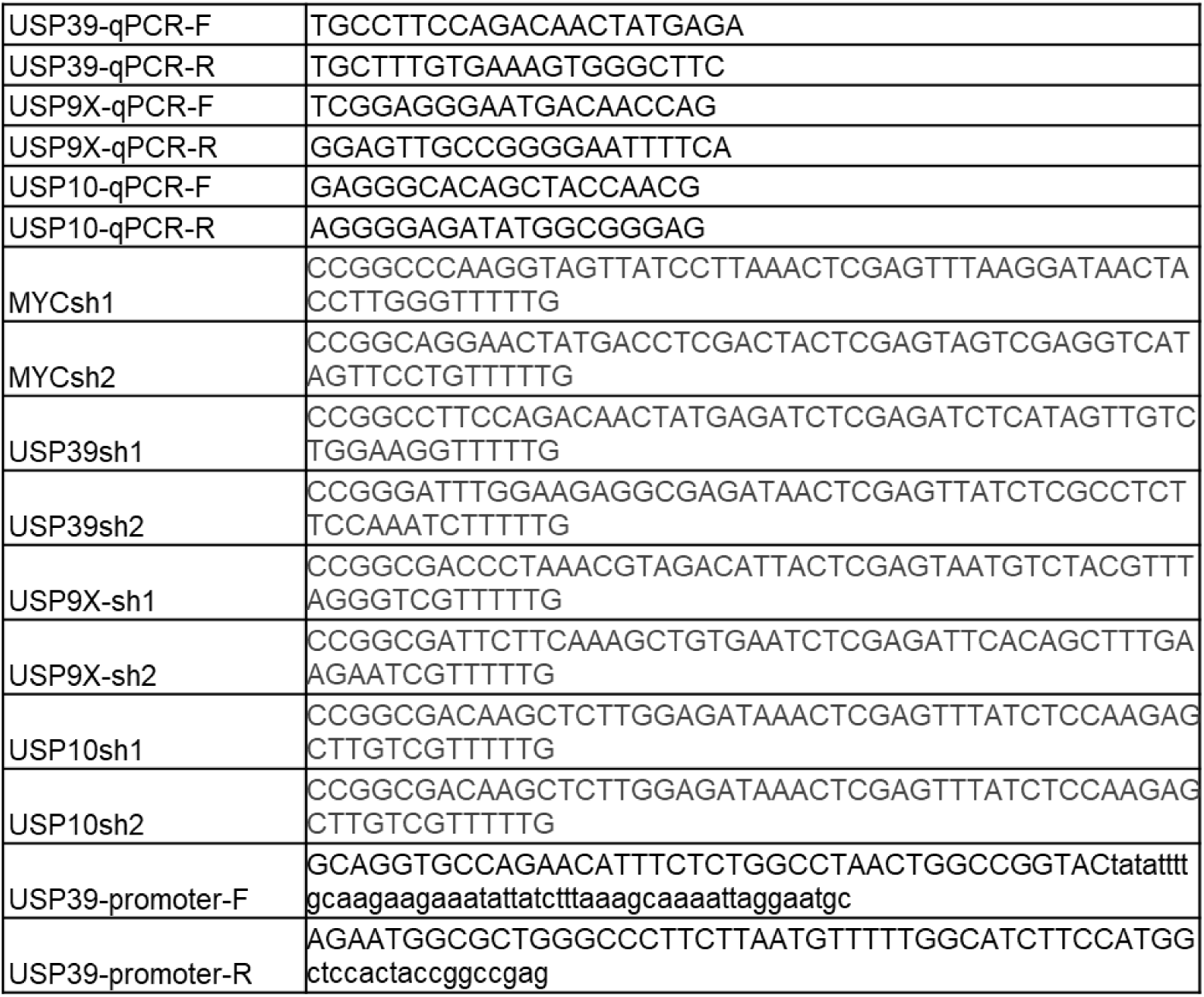
Oligonucleotides.

Among the four proteins, PSMD7 and PSMD1 are non-ATPase regulatory subunits of the 19*S* proteasome complex, which functions in recognizing and processing polyubiquitinated proteins for degradation by the 26*S* proteasome.^29^ CUL4B is a core scaffold protein of the CRL4B E3 ubiquitin ligase complex that facilitates the recruitment of substrate adaptors and promotes ubiquitin conjugation to target proteins.^30^ As an ortholog of yeast Sad1, USP39 was initially identified as a component of the spliceosome that promotes U4/U6.U5 tri-snRNP assembly and the transition of spliceosomal complex B to B^act^.^31, 32^ USP39 was recently reported as a specific DUB capable of recognizing and selectively stabilizing individual protein substrates.^33–37^ We therefore sought to further investigate the interaction between SRSF1 and USP39.

We performed immunoprecipitation in SUIT-2 and PANC-1 cells and confirmed the interaction between SRSF1 and USP39 at the endogenous protein levels (Fig. 1C and 1D). Given that SRSF1 and USP39 are both spliceosome components, we want to examine whether the interaction is mediated through RNA binding. Notably, the interaction was resistant to benzonase treatment, indicating that it is not dependent on nucleic acids (Fig. 1C and 1D). This interaction was further validated in SUIT-2 cells overexpressing SRSF1 (Fig. 1E).

SRSF1 comprises three domains: two RNA recognition motifs (RRMs) that specifically bind to RNA, and a C-terminal RS domain enriched in arginine and serine residues that mediates protein-protein interactions.^38, 39^ To identify the domain of SRSF1 responsible for interacting with USP39, we performed co-IP assays in 293T cells expressing Flag-USP39 together with various domain-deleted or mutated T7-tagged SRSF1 constructs. These included deletion mutants lacking RRM1 (ΔRRM1), RRM2 (ΔRRM2), or the RS domain (ΔRS). In addition, NRS consists of a C-terminal fusion of SRSF1 with the nuclear retention signal from SRSF2. The inactivated RRM1 mutant of SRSF1 (FFDD) has Phe56 and Phe58 substituted with Asp residues.^40, 41^ We found that the interaction was preserved in the ΔRRM1, ΔRRM2, and FFDD mutants (Fig. 1F), further supporting the finding that the interaction is RNA-independent (Fig. 1C-E). In contrast, removing the RS domain greatly weakened the interaction, indicating that this domain is critical for USP39 binding.

### USP39 stabilizes SRSF1 in a proteasome-dependent manner

Recent studies reported that USP39 could remove the conjugated ubiquitin from protein substrates and hence impair their ubiquitin-mediated degradation.^33–37^ Consistent with this mechanism, we found that USP39 knockdown decreased SRSF1 protein levels in SUIT-2 cells, which could be rescued by proteasome inhibition using MG132 (Fig. 2A). In contrast, knockdown of two other DUBs—USP9X and USP10—identified by immunoprecipitation-mass spectrometry and proximity labeling, respectively, did not significantly affect SRSF1 expression (Fig. S1).

**Figure 2.**
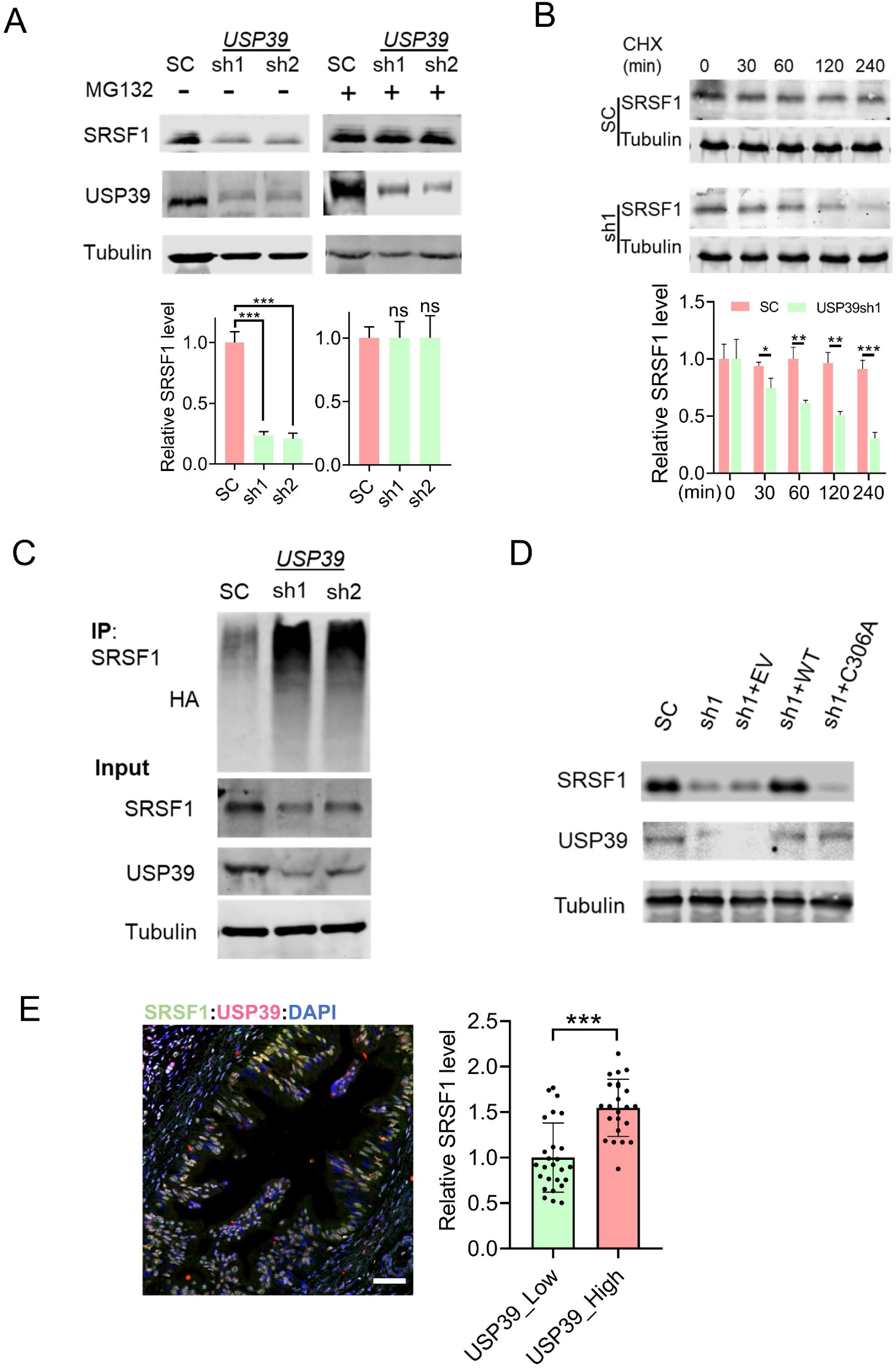
USP39 stabilizes SRSF1 by preventing its proteasomal degradation. (A) Western blot analysis of SRSF1, USP39, and Tubulin in SUIT-2 cells with *USP39* knockdown, with or without MG132 treatment. SC, scramble control. (B) Western blot analysis of SRSF1 and Tubulin in *USP39*-depleted SUIT-2 cells treated with cycloheximide (CHX; 10 µg/mL) for the indicated durations. The data are quantitated on the right (n = 3). Unpaired two-tailed t test, *p <0.05, **p <0.01, ***p <0.001. Error bars represent mean ± SD. (C) *In vivo* ubiquitination assay in SUIT-2 cells stably expressing HA–ubiquitin, with or without *USP39* knockdown. (D) Western blot analysis of SRSF1, USP39, and Tubulin in SUIT-2 cells with *USP39* knockdown, rescued by expression of either shRNA-resistant wild-type USP39 or the catalytically inactive C306A mutant. (E) IF staining (left) and quantification (right) of SRSF1 and USP39 in individual cells of a human PDAC tumor. Scale bar, 50 μm. Unpaired two-tailed t test, ***p <0.001. Error bars represent mean ± SD.

Considering that USP39 also promotes U4/U6.U5 tri-snRNP assembly and the transition of spliceosomal complex B to B^act^,^31, 32^ and that SRSF1 expression is regulated by alternative splicing within its 3’UTR,^19, 25^ we further examined whether the decrease in SRSF1 levels upon *USP39* knockdown results from defective splicing. To test this potential mechanism, we treated SUIT-2 cells with Pladienolide B, a natural macrolide compound that inhibits pre-mRNA splicing by binding to SF3B1.^42^ Interestingly, Pladienolide B treatment led to an increase in SRSF1 protein levels, while USP39 expression remained unchanged (Fig. S2). These results suggest that the downregulation of SRSF1 upon USP39 knockdown is not due to defective splicing.

To investigate whether *USP39* knockdown affects SRSF1 protein stability, we treated SUIT- 2 cells with cycloheximide (CHX) to inhibit protein synthesis. SRSF1 stability was significantly reduced following *USP39* knockdown (Fig. 2B). Consistently, *USP39* knockdown led to a marked increase in SRSF1 ubiquitination levels (Fig. 2C). Consistently, ectopic expression of shRNA-resistant USP39, but not C306A deubiquitinase-inactive mutant^43–45^, significantly rescued SRSF1 expression in SUIT-2 cells with *USP39* knockdown (Fig. 2D). We further examined the expression correlation between SRSF1 and USP39 in human PDAC tumor samples. Tumor cells with higher USP39 protein levels exhibited correspondingly elevated SRSF1 expression (Fig. 2E). These results suggest that USP39 promotes SRSF1 expression by suppressing its ubiquitination and subsequent degradation.

### Elevated USP39 expression is associated with poor prognosis in PDAC

We previously reported that SRSF1 expression is dynamically regulated during PDAC tumorigenesis and progression.^19^ Upregulation of SRSF1 promotes epithelial cell transformation and tumor development. Based on this, we examined the expression of USP39 in precancerous pancreatic lesions and PDAC samples.

Integrative analysis of datasets from TCGA (The Cancer Genome Atlas)^46^ and GTEx (Genotype-Tissue Expression project)^47^ revealed that *USP39* RNA levels are significantly elevated in PDAC tumors compared to normal pancreatic tissue (Fig. 3A). Consistently, data from published transcriptomic studies (GSE28735 and GSE16515) also indicate increased *USP39* mRNA expression in PDAC (Fig. 3B and 3C). We observed upregulation of USP39 in the KPC (*LSL-**K**ras^G12D/+^*; *LSL-Tr**p**53^R172H/+^*; *Pdx1-**C**re*) mouse model of PDAC: USP39 expression was significantly higher in PDAC tumor cells compared to normal pancreatic tissue and tumor-adjacent stroma (Fig. 3D). Notably, increased USP39 expression was already detectable in pancreatic intraepithelial neoplasia (PanIN), suggesting a potential oncogenic role of USP39 during early tumor initiation.

**Figure 3.**
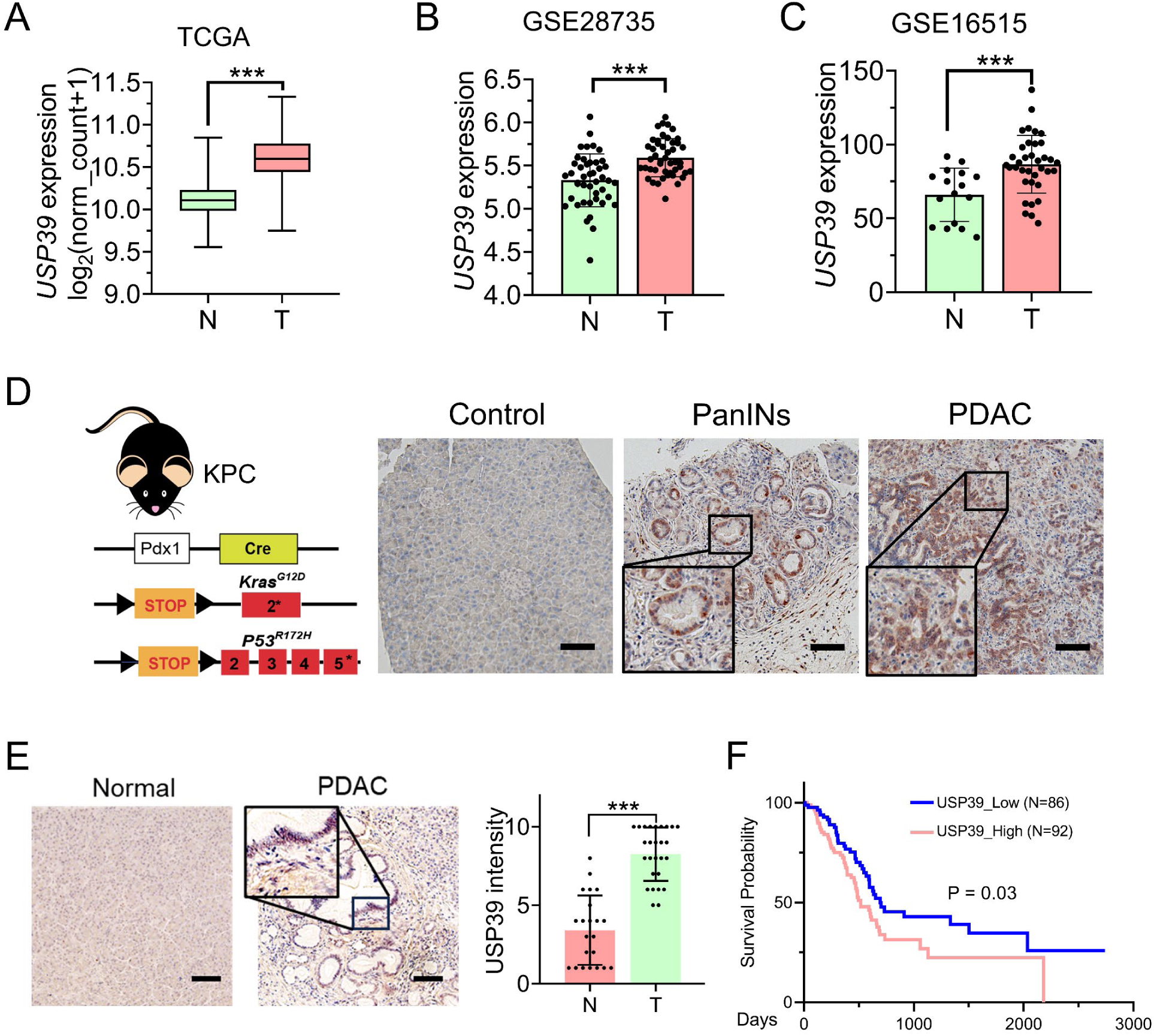
Elevated USP39 expression is associated with poor prognosis in PDAC. (A) *USP39* expression in normal pancreatic samples and PDAC tumors from TCGA RNA-sequencing data. (B and C) *USP39* expression in normal pancreatic samples and PDAC tumors based on microarray data from (B) GSE28735 and (C) GSE16515 datasets. (D) Scheme of KPC mice and IHC staining of USP39 in pancreata from control and KPC mice. Insets show higher magnification of PanIN1A lesions and PDAC tumors, respectively. Scale bar, 100 μm. (E) Representative IHC staining (left) and intensity scores (right) of USP39 in human normal pancreatic tissue (n=22) and PDAC tumors (n=27). Unpaired two-tailed t test, ***p <0.001. Error bars represent mean ± SD. (F) Kaplan–Meier survival analysis of PDAC patients stratified by *USP39* expression levels, based on the TCGA dataset.

Further evaluation of human PDAC samples revealed elevated USP39 protein levels in tumors compared to normal pancreas tissue (Fig. 3E). Importantly, patients with higher *USP39* expression exhibited significantly poorer survival outcomes than those with lower expression (Fig. 3F). These results indicate that elevated USP39 expression is associated with poor PADC prognosis.

### *USP39* knockdown suppresses PDAC progression

We next investigated the functional role of USP39 in PDAC progression. shRNA-mediated knockdown of *USP39* significantly impaired the proliferation and migration of SUIT-2 and PANC- 1 cells (Fig. 4A, 4B, and S3). These effects were fully rescued by ectopic expression of shRNA-resistant wild-type USP39, but not by the catalytically inactive C306A mutant, suggesting that the deubiquitinase activity of USP39 is critical for its oncogenic function.

**Figure 4.**
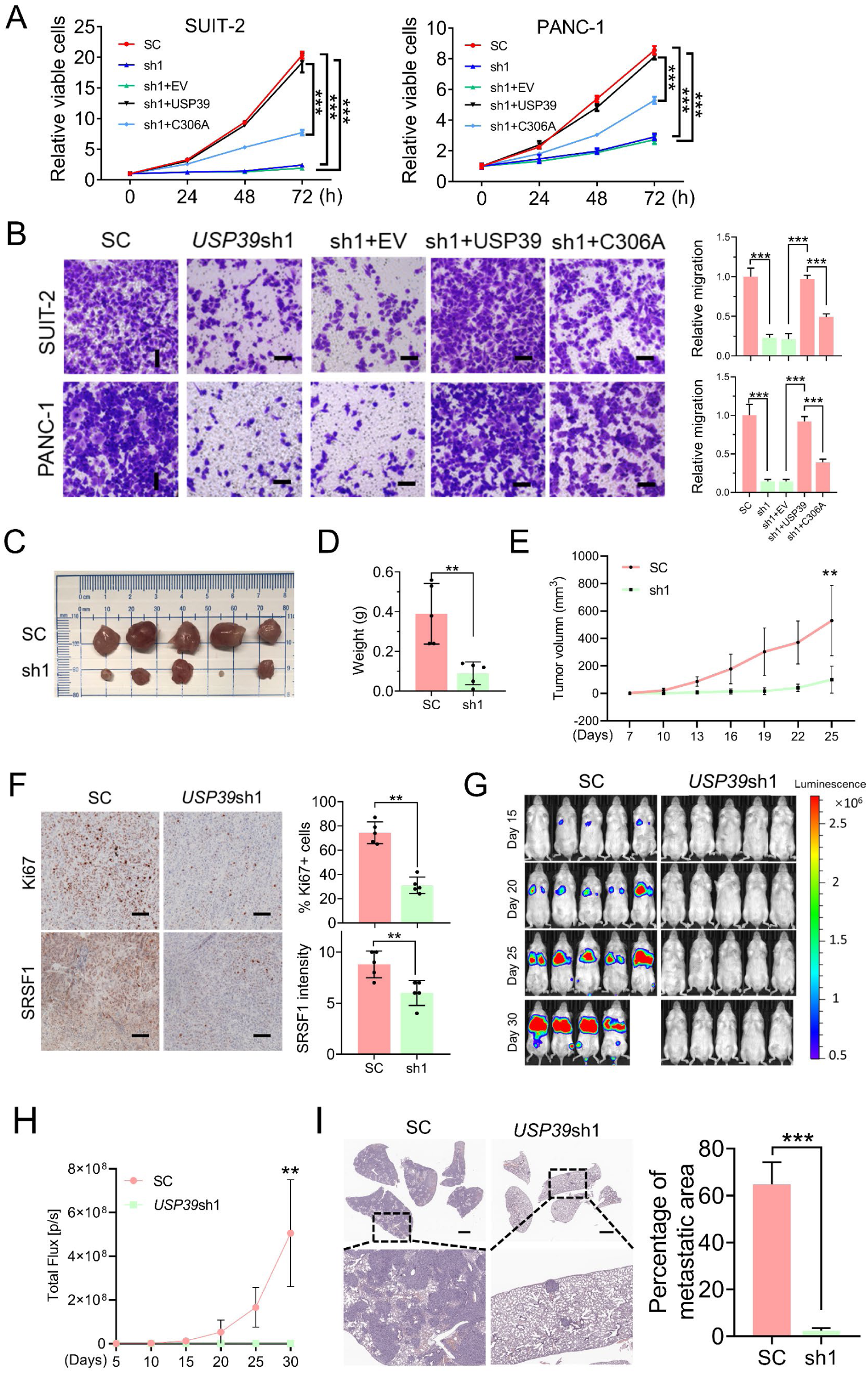
*USP39* knockdown suppresses PDAC tumor progression. (A) Proliferation of SUIT- 2 and PANC-1 cells with *USP39* knockdown, rescued by expression of either shRNA-resistant wild-type USP39 or the C306A mutant, was measured using CCK-8 assays at the indicated time points. One-way ANOVA with Tukey multiple comparisons test. ***p < 0.001; n=3. Error bars represent mean ± SD. (B) Transwell assay of the cell migration properties of SUIT-2 and PANC- 1 cells with *USP39* knockdown, rescued by expression of either shRNA-resistant wild-type USP39 or the C306A mutant; Scale bar, 150 μm. Bar graphs (right) show the quantification of relative migration. One-way ANOVA with Tukey multiple comparisons test, ***p < 0.001; n=3. Error bars represent mean ± SD. (C) Subcutaneous tumors formed by SUIT-2 cells, with or without *USP39* knockdown. (D) Weight of subcutaneous xenografts from SUIT-2 cells, with or without *USP39* knockdown. (E) Growth curve of subcutaneous tumors formed by SUIT-2 cells, with or without *USP39* knockdown. Unpaired two-tailed t-test, **p < 0.01; n = 5. Error bars represent mean ± SD. (F) IHC staining and quantification of Ki67 and SRSF1 in subcutaneous tumors formed by SUIT- 2 cells, with or without *USP39* knockdown. Scale bar, 100 μm. Unpaired two-tailed t-test, **p < 0.01; n = 5. Error bars represent mean ± SD. (G) Luciferase bioluminescence imaging of pulmonary metastatic foci following tail vein injection of SUIT-2 cells with or without USP39 knockdown. n = 5. One mouse in the control group was euthanized prior to the endpoint due to excessive tumor burden. (H) Quantification of photon flux in pulmonary metastatic luciferase foci from (G). Unpaired two-tailed t-test, **p < 0.01; Error bars represent mean ± SD. (I) H&E staining and quantification of pulmonary metastatic foci from (G). Scale bar, 2 mm. Unpaired two-tailed t- test, ***p < 0.001; Error bars represent mean ± SD.

We further investigated the oncogenic role of USP39 *in vivo*. SUIT-2 cells stably expressing *USP39* shRNA or control shRNA were subcutaneously injected into the flanks of immunodeficient nude mice to establish xenograft tumor models. Tumor growth was monitored over time, and both tumor volume and tumor weight were significantly reduced in mice bearing *USP39* knockdown tumors compared to those injected with control cells (Fig. 4C and 4D). Moreover, there was a marked suppression in the rate of tumor growth in the *USP39*-depleted group (Fig. 4E).

To further explore the effects of *USP39* depletion on tumor cell proliferation, we performed immunohistochemical analysis of Ki67. Tumors derived from *USP39* knockdown cells exhibited a notable reduction in Ki67-positive nuclei, suggesting a decrease in proliferative capacity (Fig. 4F). Immunohistochemistry also revealed a significant reduction in SRSF1 protein levels in the *USP39* knockdown tumors compared to the controls (Fig. 4F); however, this difference did not reach statistical significance when assessed by western blot analysis (Fig. S4).

To evaluate the role of USP39 in tumor metastasis, luciferase-labeled control or *USP39*- shRNA SUIT-2 cells were intravenously injected into NSG mice, and lung colonization was monitored by bioluminescent imaging (BLI). Continued BLI monitoring revealed a marked reduction in metastatic outgrowth in the *USP39* knockdown group (Fig. 4G and 4H).

Consistently, H&E staining of lung sections showed significantly fewer and smaller metastatic lesions in mice injected with USP39-depleted cells compared to controls (Figs. 4I). These findings collectively support a role for USP39 in promoting PDAC progression.

### *USP39* knockdown suppresses PDAC progression via SRSF1 downregulation

We further examined the contribution of SRSF1 regulation to the oncogenic activity of USP39. Overexpression of SRSF1 partially rescued the impaired proliferation observed in *USP39*-depleted SUIT-2 and PANC-1 cells (Fig. 5A). Consistently, *USP39* knockdown led to a marked reduction in cell division in both cell lines (Fig. 5B), an effect that was also reversed by SRSF1 overexpression. Furthermore, SRSF1 overexpression restored the migratory capacity of *USP39* knockdown cells (Fig. 5C).

**Figure 5.**
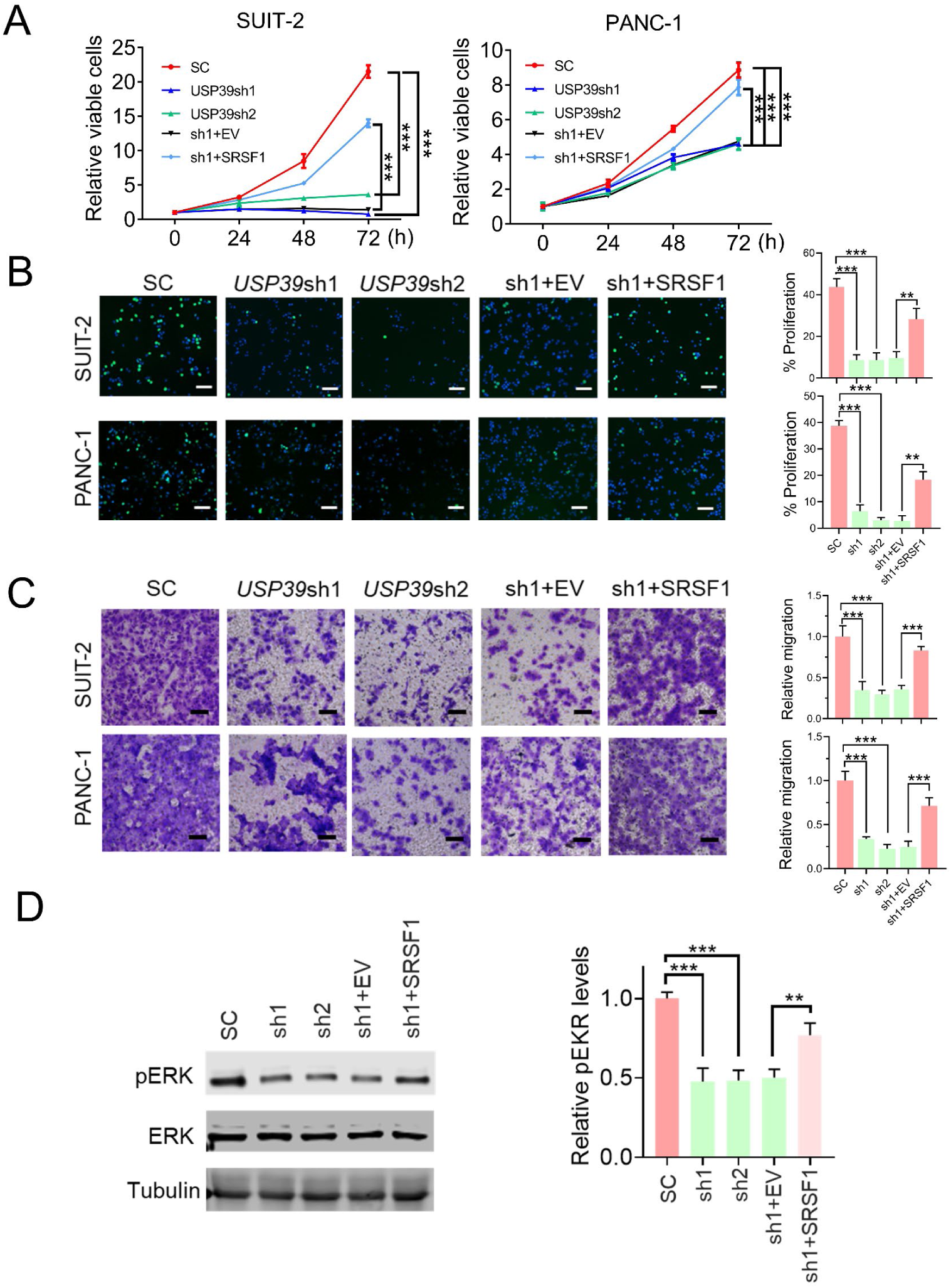
*USP39* knockdown suppresses PDAC cell proliferation and migration, at least in part through reduced SRSF1 expression. (A) The proliferation of *USP39*-knockdown SUIT-2 and PANC-1 cells with SRSF1 overexpression was measured using CCK-8 assays at the indicated time points. One-way ANOVA with Tukey multiple comparisons test. ***p < 0.001; n=3. Error bars represent mean ± SD. (B) Nuclear EdU incorporation in *USP39*-knockdown SUIT-2 and PANC-1 cells, with or without SRSF1 overexpression. The cells were incubated with EdU for 12 h; scale bar, 100 μm. Bar graphs (right) show the quantification of the percentage of proliferating cells. One-way ANOVA with Tukey multiple comparisons test, **p < 0.01, ***p <0.001; n=3. Error bars represent mean ± SD. (C) Transwell assay of the cell migration properties of *USP39*-knockdown SUIT-2 and PANC-1 cells, with or without SRSF1 overexpression; Scale bar, 150 μm. Bar graphs (right) show the quantification of relative migration. One-way ANOVA with Tukey multiple comparisons test, ***p < 0.001; n=3. Error bars represent mean ± SD. (D) Representative western blot analysis of pERK, ERK, and Tubulin in SUIT-2 cells with *USP39* knockdown, with or without SRSF1 overexpression. Quantification of pERK is shown on the right. One-way ANOVA with Tukey multiple comparisons test, **p < 0.01, ***p <0.001; n=3. Error bars represent mean ± SD.

Previous studies reported that USP39 is essential for KRAS-driven lung and colon cancers,^48^ and that its knockdown reduces phosphorylated ERK (pERK) levels in renal cell carcinoma.^49^ Given that SRSF1 plays a critical role in regulating MAPK signaling,^19^ we investigated whether USP39 influences this pathway in PDAC cells. In SUIT-2 cells, *USP39* knockdown led to a reduction in pERK, which was rescued by SRSF1 overexpression (Fig. 5D). These findings suggest that USP39 promotes PDAC progression partially by stabilizing SRSF1 and sustaining MAPK signaling.

### MYC binds to the *USP39* promoter and activates its transcription

We previously reported that hyperactive MYC overrides the negative feedback downregulation of SRSF1 in pancreatic epithelial cells expressing KRAS^G12D^.^19^ Notably, transcriptomic analysis revealed that MYC activation suppresses the ubiquitination regulation in mouse pancreas.^19^ We therefore investigated whether MYC regulates USP39 expression in PDAC. Correlation analysis of gene expression data showed a positive correlation between *MYC* and *USP39* mRNA levels in PDAC tumors (Fig. 6A). We further reanalyzed transcriptomic data from a mouse model with acute activation of quasi-physiologic levels of MYC in the pancreas, induced by tamoxifen treatment^50^ (Fig. 6B) as well as from KPC PDAC cells with doxycycline-induced *MYC* knockdown^51^ (Fig. 6C). In addition to confirming SRSF1 as a key transcriptional target of MYC^19^, we identified *USP39*—along with several other USP family members—as being upregulated following MYC activation and downregulated upon shRNA-mediated *MYC* depletion (Fig. 6C). To validate the RNA-sequencing results, we knocked down *MYC* in SUIT-2 cells and observed a significant decrease in USP39 protein levels (Fig. 6D). Consistently, *USP39* mRNA levels were also significantly reduced following *MYC* knockdown, supporting its regulation as a transcriptional target of MYC (Fig. 6E).

**Figure 6.**
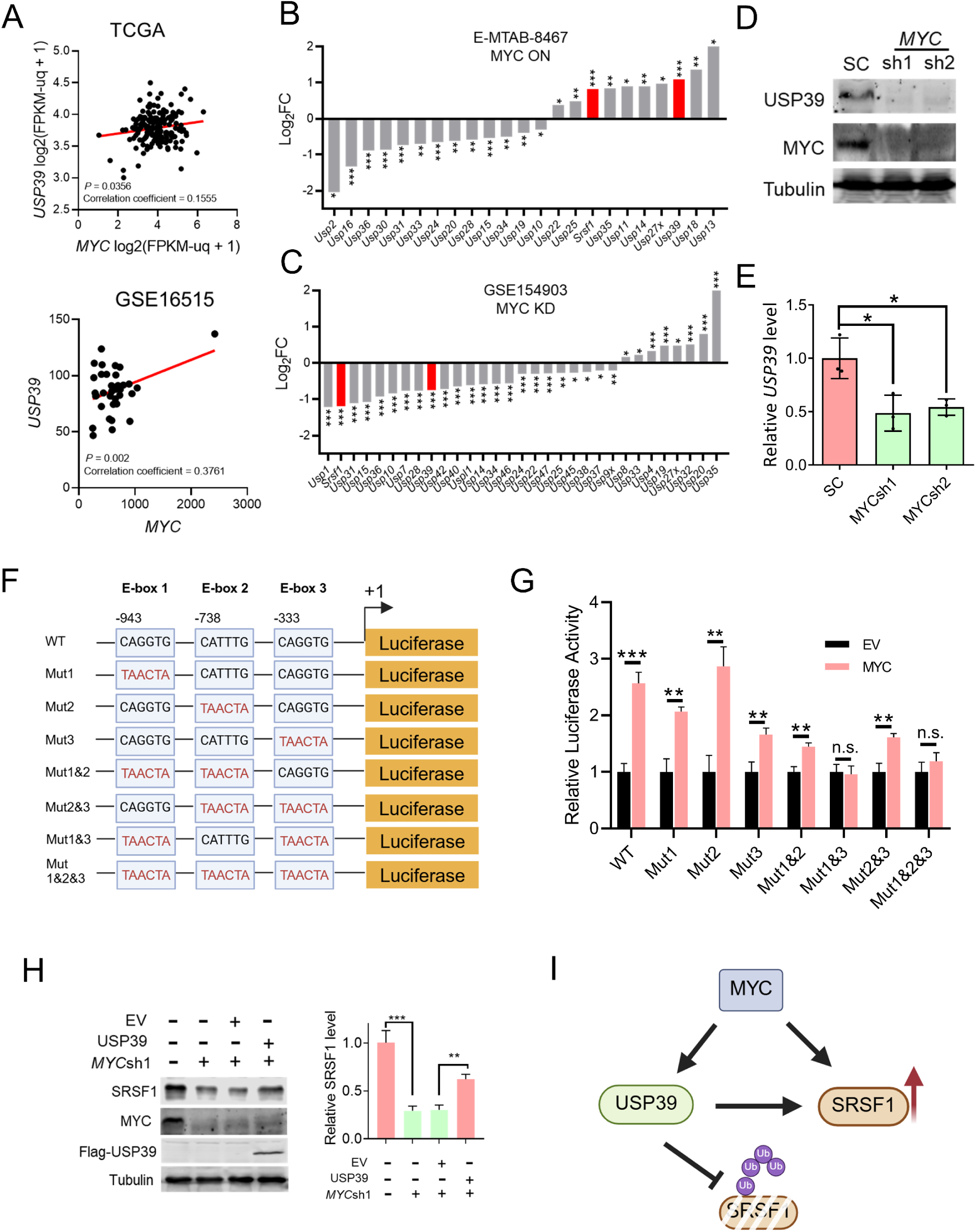
*USP39* is a transcriptional target of MYC in PDAC. (A) Correlation between *USP39* and *MYC* mRNA levels in PDAC, based on gene expression data from TCGA and GSE16515. (B) Expression changes in *Srsf1* and *Usp39* (red bars), and other USP family members, following MYC activation in the pancreas in an inducible MYC-ER mouse model. Quasi-likelihood F-test adjusted for multiple testing: *p < 0.05, **p < 0.01, ***p < 0.001. (C) Expression changes in *Srsf1* and *Usp39* (red bars) and other USP family members in KPC cells with Dox-induced knockdown of *Myc*. Quasi-likelihood F-test, adjusted for multiple testing: *p < 0.05, **p < 0.01, ***p < 0.001. (D) Western blot analysis of USP39 and MYC in SUIT-2 cells with *MYC* knockdown. (E) Real time RT-PCR of *USP39* in SUIT-2 cells with *MYC* knockdown. One-way ANOVA with Tukey multiple comparisons test, *p < 0.05; n=3. Error bars represent mean ± SD. (F) Schematic diagram of the *USP39* promoter (−1500-0) comprising three non-canonical E-boxes, and the E-Box mutants. Mutant E-boxes and residues are indicated in red. (G) Luciferase assay of reporter constructs in SUIT-2 cells co-transfected with a Renilla control plasmid, with or without MYC overexpression. Luciferase activity was normalized to Renilla luciferase. Unpaired two-tailed t test, *p < 0.05, **p < 0.01, ***p < 0.001; n=3. Error bars represent mean ± SD. (H) Western blot analysis of SRSF1, Flag-USP39, and MYC in *MYC* knockdown SUIT-2 cells, with or without USP39 overexpression. (I) MYC forms a regulatory network by increasing USP39 and SRSF1 expression, and USP39 stabilizes SRSF1 by decreasing its ubiquitination.

We then investigated whether MYC directly binds to the *USP39* promoter. The *USP39* gene contains two alternative transcriptional start sites (Fig. S5A). ChIP-seq occupancy profiles of MYC, based on data from K562, HeLa-S3, HepG2, and HUVEC cells, indicate MYC binding at both *USP39* promoters. Notably, analysis of TCGA RNA-sequencing data suggests that only exon 1b is actively transcribed in pancreatic cancer (Fig. S5B). We therefore focused on MYC-mediated regulation of the *USP39* promoter associated with exon 1b.

Three putative MYC binding sites (E-box motifs^52–55^) were identified within this promoter region. We generated a luciferase reporter construct containing the 1.5 kb region upstream of *USP39* exon 1b (Fig. 6F). MYC overexpression resulted in a two-fold increase in luciferase activity, indicating transcriptional activation. To determine which MYC binding sites contribute to this regulation, we introduced mutations into each E-box motif individually and in various combinations (Fig. 6F). Mutation of E-box2 had no significant effect on MYC-mediated transcriptional activation. In contrast, mutations in E-box1 or E-box3 partially reduced luciferase activity. Notably, combined mutations in E-box1 and E-box3, or in all three E-boxes, completely abolished MYC-induced transcriptional activation (Fig. 6G). These results suggest that MYC directly regulates *USP39* transcription through cooperative binding at multiple E-box elements within the exon 1b promoter.

Given that hyperactive MYC promotes PDAC progression in part through increased SRSF1 expression^19, 56^, we investigated whether USP39 contributes to this regulatory axis. We found that MYC knockdown reduced SRSF1 expression, whereas USP39 overexpression partially restored SRSF1 levels (Fig. 6H). These findings suggest that USP39 acts downstream of MYC to support SRSF1 expression, potentially linking MYC-driven transcriptional programs to post-translational stabilization mechanisms in PDAC (Fig. 6I).

## DISCUSSION

Aberrant RNA splicing drives tumorigenesis through the misregulation of diverse signaling pathways.^13–16^ As a key proto-oncogenic splicing factor,^25, 56–60^ SRSF1 regulates RNA splicing during the early stages of spliceosome assembly, in part by facilitating splice-site selection through the recruitment of core spliceosomal proteins.^17^ SRSF1 activity is regulated not only at the transcriptional level, but also through various post-transcriptional mechanisms—including alternative splicing coupled with nonsense-mediated mRNA decay (NMD), mRNA nuclear retention, translational repression, and regulation by microRNAs such as miR-7—as well as through multiple post-translational modifications.^20–25, 28, 61^

In early-stage pancreatic cells, SRSF1 is destabilized through increased ubiquitination as a negative feedback mechanism in response to KRAS^G12D^ expression.^19^ This post-translational regulation acts to constrain MAPK signaling and limit oncogenic transformation. However, transcriptional activation of *SRSF1* by MYC overrides this homeostatic feedback mechanism, resulting in elevated SRSF1 and promoting PDAC progression. Here, we further demonstrate that, in addition to transcriptional activation, MYC enhances SRSF1 protein stability through the deubiquitinase USP39. MYC directly binds to the *USP39* promoter and upregulates its transcription. In turn, USP39 reduces the ubiquitination levels of SRSF1 by interacting with its RS domain in PDAC cells. Notably, USP39 expression is already elevated in precancerous pancreatic lesions, suggesting its potential involvement in the early stages of tumor initiation and progression. We previously reported that SRSF1 is upregulated and promotes pancreatitis.^19^ Further investigation is warranted to understand the role of USP39 during early cellular transformation events, such as acinar-to-ductal metaplasia, and in the context of pancreatitis.

Multiple studies have identified USP39 as an essential component of the spliceosome, where it stabilizes the interaction between U4/U6 and U5 snRNPs during spliceosome assembly^31, 32, 62^ Rather than recapitulating the decrease in SRSF1 levels observed with *USP39* knockdown, we found that pharmacological inhibition of RNA splicing using Pladienolide B led to an increase in SRSF1 expression, suggesting a compensatory response to splicing disruption.

Together with our observation that the interaction between USP39 and SRSF1 is RNA-independent, these findings suggest that USP39 regulates SRSF1 expression through a mechanism distinct from its canonical spliceosomal function. Nonetheless, it remains possible that their interaction is facilitated by spatial proximity within the spliceosome.

Our findings demonstrate that USP39 functions as an oncogenic factor in PDAC. USP39 expression is significantly elevated in both human PDAC tumors and early precursor lesions, with higher levels correlating with poor patient survival. Functional studies reveal that USP39 promotes PDAC cell proliferation and migration, and supports tumor growth *in vivo*. Moreover, our data show that SRSF1 overexpression only partially rescues the suppression of MAPK signaling and malignant phenotypes observed upon *USP39* knockdown, indicating that USP39 likely contributes to PDAC progression through additional mechanisms. These may include the deubiquitination of other substrates^34, 36, 37^ and/or its canonical role in splicing regulation^31, 32, 48^, highlighting the multifaceted functions of USP39 in tumor biology. In particular, depending on the cellular context and tissue type, specific DUB can function oppositely as tumor suppressors or oncogenes.^63, 64^ Future studies should focus on the development and comprehensive evaluation of USP39 inhibitors as potential therapeutic agents in PDAC.

In summary, we identified a MYC–USP39–SRSF1 regulatory axis that integrates both transcriptional and post-translational control mechanisms in PDAC. MYC transcriptionally activates the expression of both *USP39* and *SRSF1*; in addition, USP39 stabilizes SRSF1 protein by suppressing its ubiquitination, thereby amplifying SRSF1’s oncogenic splicing activity. These findings highlight USP39 as a potential therapeutic target in pancreatic cancer.

## Materials and Methods

### Analysis of Public Datasets

mRNA expression levels of *USP39*, *SRSF1*, and *MYC*, along with relevant clinical information, were retrieved from the Gene Expression Omnibus (GEO), The Cancer Genome Atlas (TCGA), and the Genotype-Tissue Expression (GTEx) databases. Expression comparisons between pancreatic tumor and normal tissues were performed using GEO datasets GSE16515 and GSE28735, as well as the UCSC Xena platform^65^ incorporating TCGA and GTEx data. To analyze transcriptomic changes in response to MYC modulation, we retrieved public RNA-seq data obtained from a transgenic mouse model with acute pancreatic MYC activation^50^ and from *MYC* knockdown in KPC cells^51^. Raw read counts for annotated genes were generated using featureCounts^66^, and differential gene expression analysis was performed using edgeR. All sequencing data analyses were conducted on the SeaWulf high-performance computing cluster at Stony Brook University.

Publicly available MYC ChIP-seq datasets were analyzed using the UCSC Genome Browser (https://genome.ucsc.edu). ChIP-seq tracks for MYC binding were retrieved from ENCODE datasets for human cell lines, including K562, HeLa-S3, HepG2, and HUVEC.

### Statistical Analysis

Statistical analyses were performed using GraphPad Prism and R. The specific statistical tests used for each experiment are detailed in the corresponding figure legends. All graphs display individual data points to indicate n values for each treatment or genotype group, with horizontal bars representing the mean, and error bars indicating the standard deviation (SD). Statistical significance is reported as follows: *, P < 0.05; **, P < 0.01; ***, P < 0.001.

### Tissue Culture

SUIT-2, PANC-1, and HEK293T cells were cultured in Dulbecco’s Modified Eagle Medium (DMEM; Thermo Fisher Scientific) supplemented with 10% fetal bovine serum (FBS; Sigma-Aldrich) and 1% penicillin-streptomycin (Thermo Fisher Scientific). Cells were maintained at 37 °C in a humidified incubator with 5% CO₂.

### Western Blotting

Cells were harvested and lysed in RIPA buffer (50 mM Tris-HCl, pH 7.4; 150 mM NaCl; 0.1% SDS; 1% NP-40; 0.5% sodium deoxycholate; 1 mM PMSF; and protease inhibitor cocktail [Roche]) by incubating on ice for 30 min. For protein stability and half-life analysis, cells were treated with 10 μg/mL cycloheximide (Sigma-Aldrich) for the indicated time points. Western blotting was performed using the following primary antibodies at 1:1000 dilution: rabbit anti- SRSF1 (Abcam), rabbit anti-USP39 (Cell Signaling Technology), rabbit anti-Tubulin (Proteintech), rabbit anti-T7 (Proteintech), rabbit anti-ERK (Cell Signaling Technology), rabbit anti-phospho-ERK (Cell Signaling Technology), rabbit anti-MYC (Cell Signaling Technology), rabbit anti-HA (Proteintech), and mouse anti-Flag (MilliporeSigma). Blots were imaged and quantified using the Odyssey Infrared Imaging System (LI-COR).

### RNA Extraction, RT-PCR, and RT-qPCR

Total RNA was extracted using TRIzol reagent (Thermo Fisher Scientific) according to the manufacturer’s instructions. Reverse transcription was performed using oligo(dT)18 primers and ImProm-II Reverse Transcriptase (Promega). PCR products were separated on agarose gels and visualized using an Azure 300 imaging system (Azure Biosystems). For RT-qPCR, cDNA was analyzed on a QuantStudio 3 Real-Time PCR System (Thermo Fisher Scientific). Relative gene expression levels were calculated using the ΔΔCq method. Primer sequences are provided in Supplementary Table 2.

### Dual-Luciferase Assay

A 1.5 kb fragment upstream of USP39 exon 1b was amplified and cloned into the pGL4.53 luciferase reporter vector (Promega) using Gibson Assembly (New England Biolabs). Dual-luciferase assays were performed using the Dual-Luciferase Reporter Assay System (Promega) according to the manufacturer’s protocol. 2 × 10^5^ SUIT-2 cells were seeded in 12-well plates.

After 24 hours, cells were co-transfected with the firefly luciferase reporter construct (300 ng) and the pRL-SV40 Renilla luciferase control vector (230 ng) using Lipofectamine 3000 (Invitrogen). Luminescence from Firefly and Renilla luciferase was measured 24 h post-transfection using a SpectraMax M5 plate reader (Molecular Devices).

### Lentiviral Generation and Infection

Lentivirus was produced in HEK293T cells by co-transfecting the expression plasmid with packaging plasmids pMD2.G and psPAX2 using Lipofectamine™ 2000 transfection reagent (Thermo Fisher Scientific). HEK293T cells were seeded one day prior to transfection at 70–80% confluency in 10-cm tissue culture plates. On the day of transfection, the medium was replaced with 9.6 mL of antibiotic-free DMEM supplemented with 10% fetal bovine serum (FBS). For each plate, 10 μg of total plasmid DNA was mixed with Lipofectamine 2000 in 400 μL of DMEM, incubated for 15 min at room temperature, and then added to the cells. After 24 h, the medium was replaced with fresh DMEM containing 10% FBS. Lentivirus-containing supernatant was collected at 48 and 72 h post-transfection, then filtered through a 0.45 μm syringe filter. Target cells were infected with viral supernatant, and 48 h post-infection, puromycin (2 μg/mL; Sigma-Aldrich) was added for selection.

### In Vivo Ubiquitination Assay

Stable HA-ubiquitin–expressing SUIT-2 cells were infected with either scramble control or USP39 short hairpin RNAs, as described above. Cells were harvested and lysed in IP lysis buffer (25 mM Tris-HCl, pH 7.5; 200 mM NaCl; 0.5% NP-40; 2 mM MgCl₂; 2 mM EDTA; 100 mM NaF) supplemented with one EDTA-free protease inhibitor tablet (Sigma-Aldrich) per 50 mL and 25 U/mL Benzonase (Sigma-Aldrich). Lysates were incubated on ice for 10 min, cleared by centrifugation, and protein concentrations were quantified and normalized using the Bradford assay. Equal amounts of protein were incubated with anti-SRSF1 antibody and Protein A/G magnetic beads (Thermo Fisher Scientific) for 3 h at 4 °C.

### Immunohistochemistry and Immunofluorescence

Tissue sections were deparaffinized, rehydrated, and treated with 3% hydrogen peroxide in methanol for 15 min to block endogenous peroxidase activity. Antigen retrieval was performed by boiling slides in 10 mM sodium citrate buffer containing 0.05% Tween-20 for 5 min. For IF analysis, slides were blocked with peptide blocking solution (Innovex Biosciences, Richmond, CA, USA) for 30 min and incubated overnight at 4 °C with mouse anti-SRSF1 (1:1000; Thermo Fisher Scientific) and rabbit anti-USP39 (1:1000; Cell Signaling Technology) antibodies. Alexa Fluor 488- or 555-conjugated, species-specific cross-adsorbed secondary antibodies (Thermo Fisher Scientific) were used for detection. Slides were mounted using ProLong Gold Antifade Mountant (Thermo Fisher Scientific). For IHC analysis, slides were incubated with HRP-labeled anti-rabbit polymer and visualized using DAB (3,3’-diaminobenzidine; DAKO). Counterstaining was performed with hematoxylin (Sigma-Aldrich), and slides were mounted using limonene-based mounting medium (Abcam).

### CCK-8 Cell Proliferation Assay

1,000 cells were seeded per well in 96-well plates in 100 µL of culture medium. Cell viability was assessed using a Cell Counting Kit-8 (CCK-8; MedChemExpress) according to the manufacturer’s instructions. Briefly, 10 µL of CCK-8 reagent was added to each well and incubated for 3 h at 37 °C. Absorbance was measured at 450 nm using a microplate reader to quantify viable cells. Each condition was measured in triplicate.

### EdU Cell Proliferation Assay

5-ethynyl-2’-deoxyuridine (EdU) incorporation was detected using an EdU-Click Chemistry 488 Kit (Sigma-Aldrich) following the manufacturer’s instructions. Briefly, cells were pulsed with 10 μM EdU. After treatment, cells were fixed, washed with PBS, and blocked as described above. They were then incubated at room temperature for 30 min with the click chemistry reaction cocktail. Following incubation, cells were washed three times with PBS. Subsequent processing was carried out according to the IF protocol described above.

### Transwell Migration Assays

Cell motility was assessed using Transwell chamber plates (24-well format, 8-μm pore size; Corning Costar). Cells that migrated to the lower surface of the membrane were fixed, stained, and imaged at 10× magnification. To quantify migration, the area covered by cells on the lower membrane surface was measured using ImageJ software (National Institutes of Health).

### Subcutaneous Tumor Growth in Nude Mice

All animal protocols were approved by the Institutional Animal Care and Use Committee at Stony Brook University. For subcutaneous tumor implantation, 10^6^ cells were resuspended in 100 μL of 1× PBS and injected into each flank of 6-week-old male BALB/c nu/nu nude mice. Tumor growth was monitored every 3 d starting from day 7 post-injection, using caliper measurements. Tumor volume was calculated using the formula: length × width² × 0.5. All mice were euthanized on day 25. Harvested tumors were bisected; one half was snap-frozen in liquid nitrogen and stored at −80 °C, and the other half was fixed in 10% neutral-buffered formalin, embedded in paraffin, sectioned, and stained for histological analysis.

### Mouse Tail-vein Assay

SUIT-2 cells stably expressing luciferase were treated with scramble control (SC) or *USP39* shRNA. 1 × 10^5^ viable SUIT-2 cells suspended in 100 μL PBS were injected into the tail vein of NSG mice. Bioluminescence imaging was initiated 5 days post-injection. Mice were first injected with luciferin (300 mg/kg, 5 min prior to imaging), anesthetized with 3% isoflurane, and then imaged in an IVIS spectrum imaging system (Caliper).

### Immunoprecipitations

Antibody binding and crosslinking to Protein A/G magnetic beads (Thermo Fisher Scientific) were performed according to the manufacturer’s instructions. Cells were harvested and lysed in IP lysis buffer. Lysates were then incubated with 2 μg of anti-SRSF1 antibody (rabbit polyclonal, Abcam or Thermo Fisher Scientific) and 30 μL of Protein A/G magnetic beads per sample.

Immunoprecipitations were carried out with rotation for 45 min at 4 °C, followed by five washes with lysis buffer to remove non-specific interactions. Bound proteins were then eluted and analyzed by SDS-PAGE and western blotting.

### Mass Spectrometry

Liquid chromatography–tandem mass spectrometry (LC-MS/MS) analysis was performed using a Fusion Lumos Tribrid mass spectrometer (Thermo Fisher Scientific) equipped with a nano-electrospray EasySpray ion source. Precursor ions were acquired every 2 sec at a resolution of 60,000 using the Orbitrap mass analyzer (m/z range: 350–1600; AGC target: 1 × 10^6^; maximum injection time: 50 ms; RF lens: 30%). Data-dependent MS/MS acquisition was performed with selection restricted to monoisotopic precursor ions with charge states 2–7, intensity >10,000, and dynamic exclusion of previously fragmented ions for 10 sec (±10 ppm tolerance). Selected precursors were isolated using a quadrupole (Q1) isolation window of 1.2 Th and fragmented by higher-energy collisional dissociation (HCD) with a normalized collision energy of 33% using the Top Speed algorithm. Fragment ions were detected in centroid mode using the linear ion trap in “Rapid” scan mode (AGC target: 10,000; maximum injection time: 35 ms). Raw data files were analyzed using the Mascot search engine against the UniProt human reference proteome database (release: September 10, 2018). MS1 mass tolerance was set to 30 ppm and MS2 tolerance to 0.2 Da. Searches assumed trypsin/P cleavage specificity with up to two missed cleavage sites. Carbamidomethylation (carboxyethylthiolation) of cysteines was set as a fixed modification, and oxidation of methionine residues was specified as a variable modification.

## Acknowledgments

We gratefully acknowledge the support of the Stony Brook University (SBU) Division of Laboratory Animal Resources, Biobank, Microscopy, and Histology Shared Resources, as well as the Mass Spectrometry Shared Resource at Cold Spring Harbor Laboratory (CSHL). Funding: L.W. was supported in part by Institutional Research Grant 21–143-01-IRG from the American Cancer Society. A.R.K. and A.J.K. were supported by NCI Program Project Grant CA13106 (Project 2). D.A.T. was supported by NIH grant R01 CA249002 and the Lustgarten Foundation. Y.P. was supported by NCI grant R50 CA211506. CSHL Shared Resources used in this study were funded in part by the NCI Cancer Center Support Grant 5P30CA045508.

## Author Contributions

Conceptualization: B.M., L.W.; Methodology: B.M., A.J.K., X.Z., A.D., P.C., Y.P., D.A.T., A.R.K., L.W.; Investigation: B.M., A.J.K., X.Z., N.S., A.D., Y.P., L.W.; Writing: B.M., A.K., P.C., Y.P., D.A.T., A.R.K., L.W.; Resources: Y.P., D.A.T., A.R.K., L.W.; Supervision: L.W.

---

**Figure S1.**
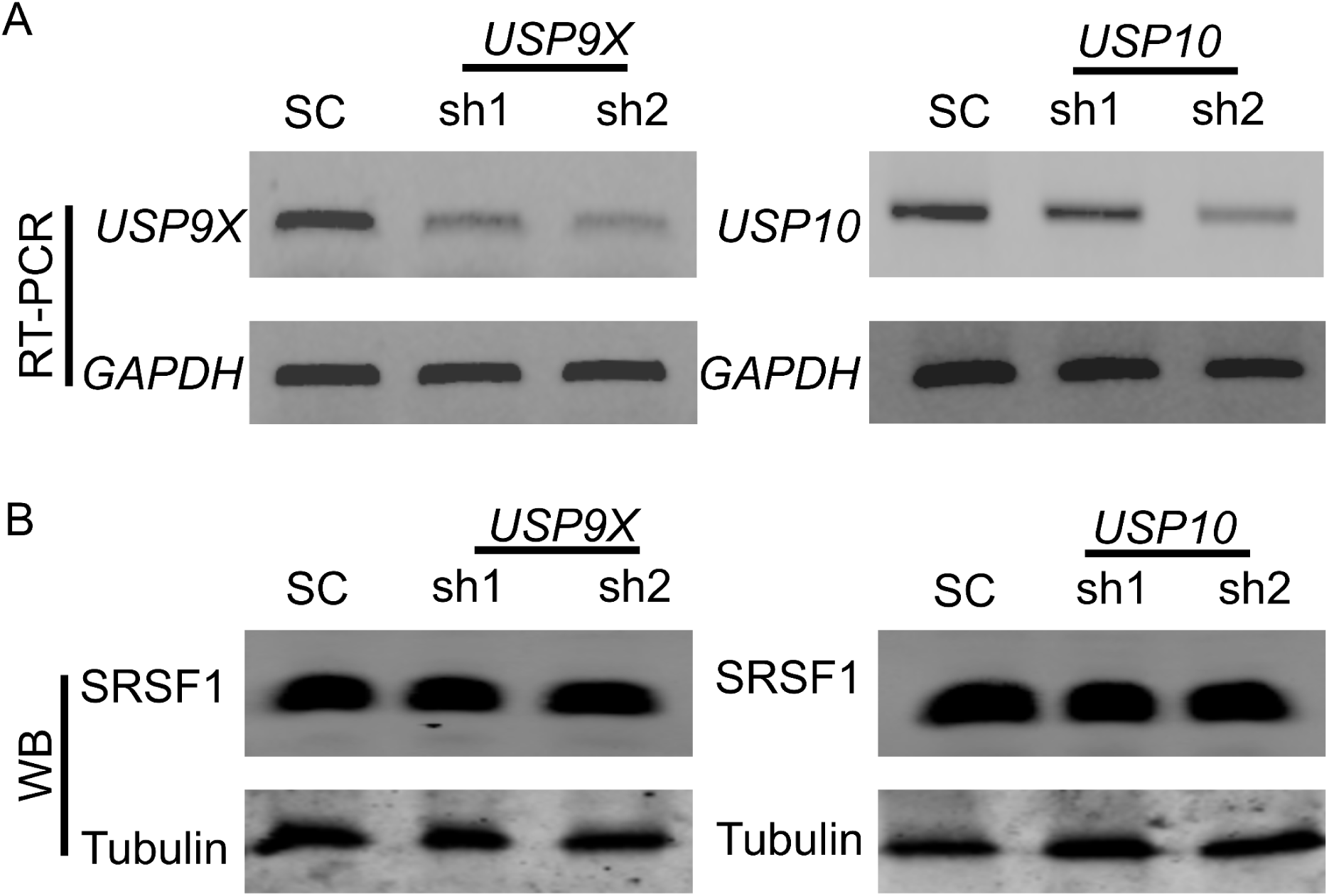
SRSF1 protein levels remain unchanged following knockdown of *USP9X* or *USP10*. (A) RT-PCR of *USP9X* and *USP10*, and (B) Western blot analysis of SRSF1 and Tubulin in SUIT-2 cells with *USP9X* and *USP10* knockdown, respectively. SC, scramble control.

**Figure S2.**
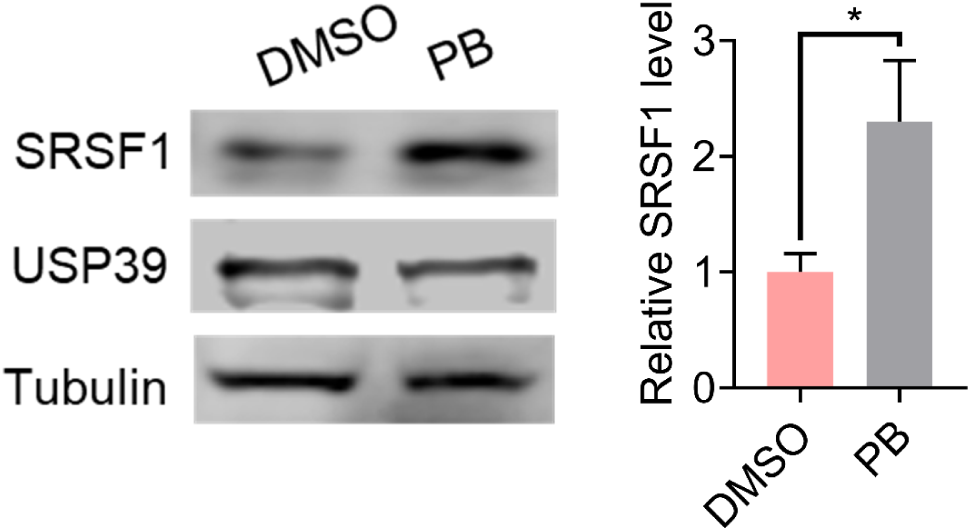
SRSF1 expression is increased upon pharmacological inhibition of RNA splicing. Western blot analysis of SRSF1, USP39, and Tubulin in SUIT-2 cells treated with 1 nM Pladienolide B (PB). Quantification of SRSF1 is shown on the right. Unpaired two-tailed t test, *p < 0.05; n=3. Error bars represent mean ± SD.

**Figure S3.**
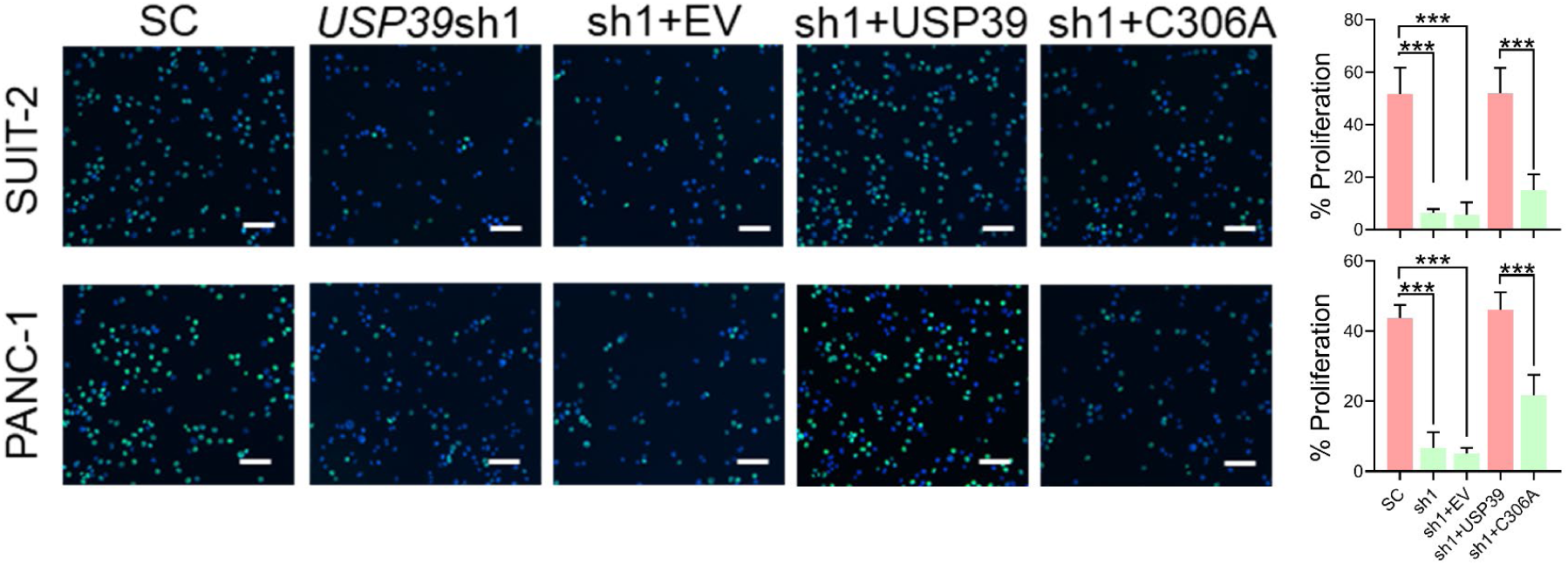
*USP39* knockdown suppresses PDAC tumor cells division. Nuclear EdU incorporation in *USP39*-knockdown SUIT-2 and PANC-1 cells, rescued by expression of either shRNA-resistant wild-type USP39 or the C306A mutant. The cells were incubated with EdU for 12 h; scale bar, 100 μm. Bar graphs (right) show the quantification of the percentage of proliferating cells. One-way ANOVA with Tukey multiple comparisons test, ***p <0.001; n=3. Error bars represent mean ± SD.

**Figure S4.**
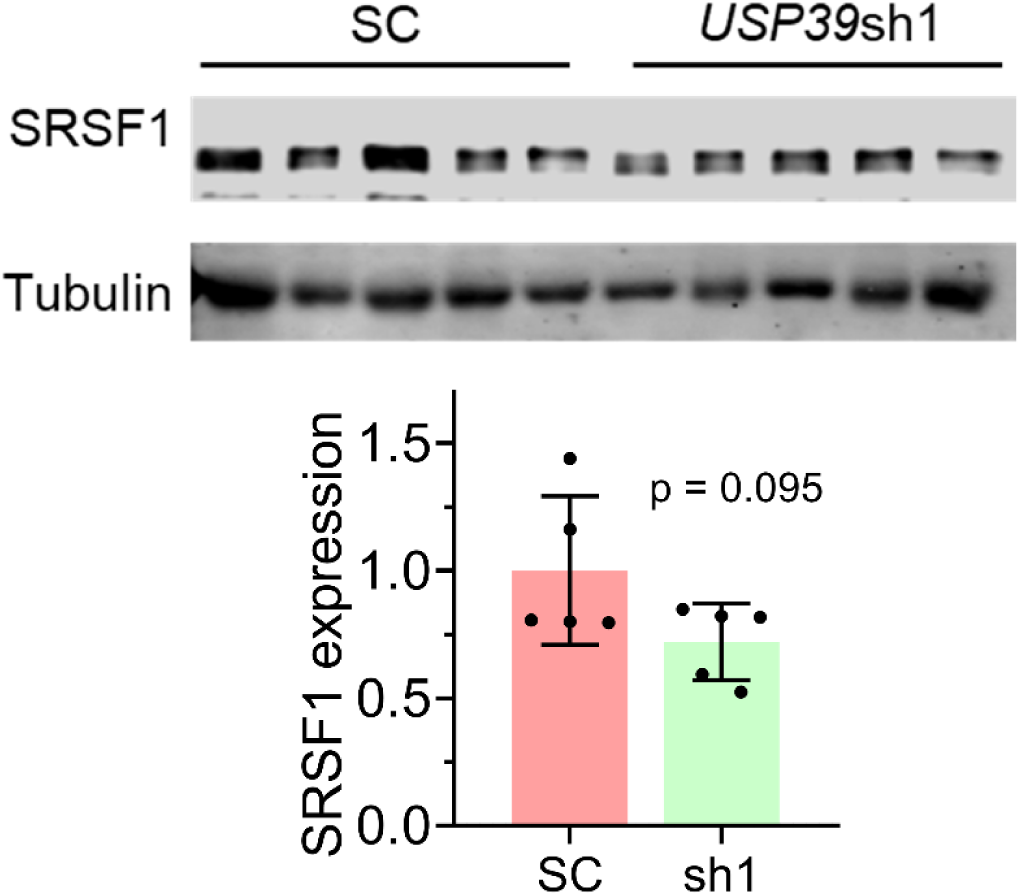
SRSF1 expression in subcutaneous xenografts, following *USP39* knockdown. Western blot analysis and quantification of SRSF1 in subcutaneous tumors formed by SUIT-2 cells, with or without *USP39* knockdown. Quantification of SRSF1 is shown on the right. Unpaired two-tailed t test. Error bars represent mean ± SD.

**Figure S5.**
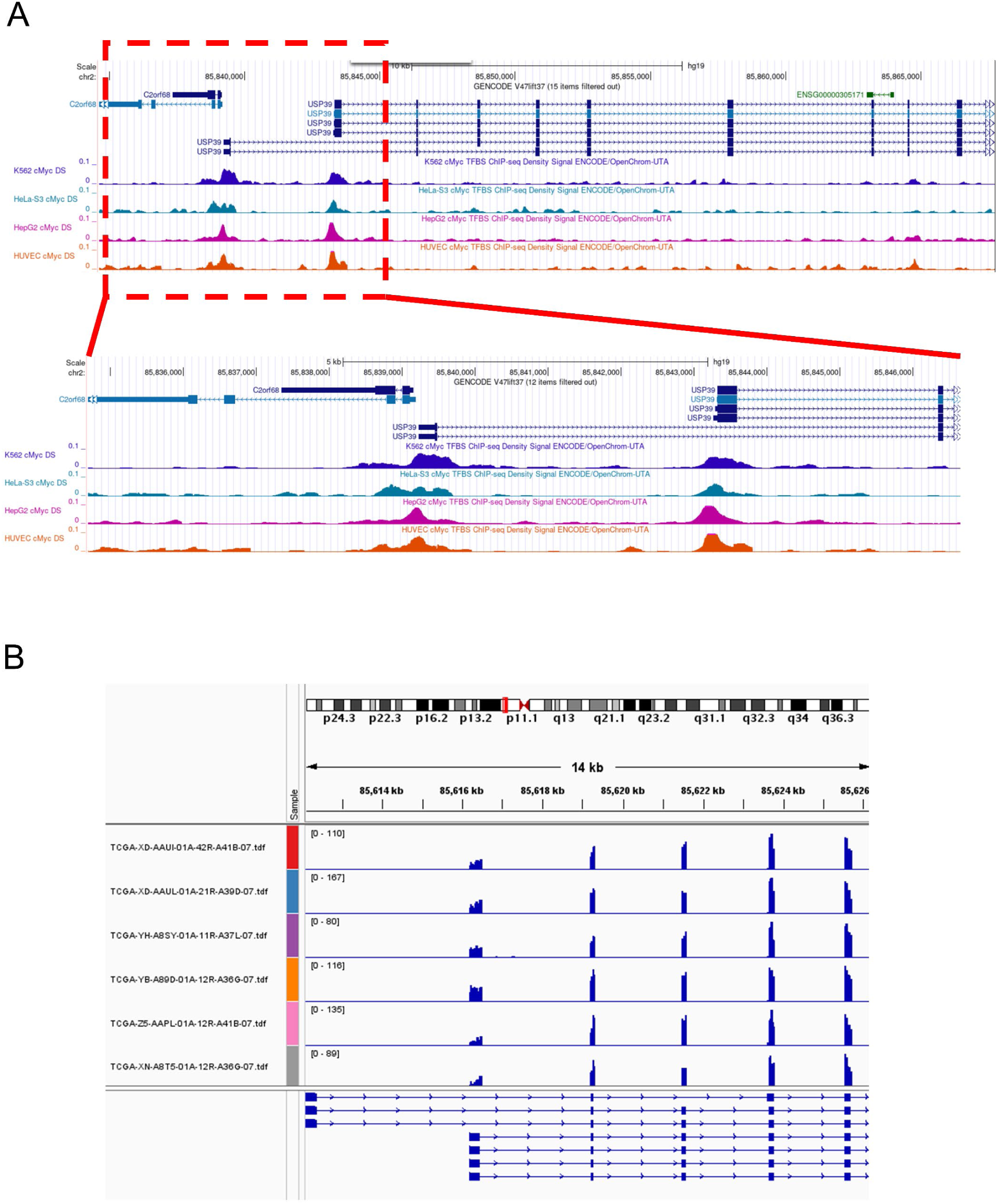
MYC binding to the promoter region of *USP39*. (A) ChIP-seq occupancy profiles of MYC at the *USP39* promoter, based on data from K562, HeLa-S3, HepG2, and HUVEC cells, sourced from the ENCODE project. (B) Representative alignment of TCGA RNA-sequencing data from pancreatic cancer samples, mapped to the *USP39* gene.

